# Stc2a inhibits IGF-stimulated somatic growth in favor of organismal survival under hypoxia

**DOI:** 10.1101/2025.10.16.682858

**Authors:** Zhengyi Wang, Jinay Shah, Shuang Li, Shriya Jaggi, Hui Xu, Cunming Duan

**Affiliations:** Department of Molecular, Cellular, and Developmental Biology, University of Michigan, Ann Arbor, MI48109

**Keywords:** Stanniocalcin 2, Insulin-like growth factor, IGF1 receptor, Papp-a, Hypoxia-inducible factor, Zebrafish

## Abstract

In response to hypoxia, animals reduce somatic growth to shift energy resources toward the maintenance of vital functions and survival. Although this phenomenon is widespread in the animal kingdom, the factors and mechanisms involved remain poorly understood. Here we report that hypoxia causes major changes in zebrafish transcriptomic landscapes with hormonal activity or hormonal signaling identified as most prominently up-regulated GO term and KEGG pathway. Among the top in this group is Stanniocalcin 2a (Stc2a), a secreted glycoprotein that inhibits insulin-like growth factor (IGF) signaling by binding to pappalysin metalloproteinases and inhibiting their activities. The hypoxic induction of *stc2a* expression is attenuated in Hif2-deficient fish. Genetic deletion of Stc2a increased the developmental speed and growth rate, resulting in enlarged adult organ and body size. Under hypoxia, *stc2a*^-/-^ fish grew faster than wild-type fish but showed reduced survival rate. These phenotypes were reversed by inhibiting pappalysin activity or blocking IGF signaling. These findings suggest that Stc2a limits IGF-mediated growth in favor of survival and that the induction of Stc2a is part of a conserved mechanism regulating the trade-off between somatic growth and survival under hypoxic stress.

## Introduction

Hypoxia poses a fundamental bioenergetic challenge to animals and their cells. Hypoxia has been linked to human diseases, such as ischemia, inflammation, tumorigenesis, and intrauterine growth restriction. In response to low oxygen levels, cells shift their metabolism toward glycolysis and reduce overall metabolic demands by altering gene expression, primarily through the actions of hypoxia-inducible factors (HIFs) (Semenza, 2012). HIFs, including HIF1, HIF2, and HIF3, are dimeric proteins composed of an oxygen-regulated α subunit and a constitutive β subunit (Semenza, 2012; Duan, 2016). Under normoxic conditions, HIF-α subunits are hydroxylated by prolyl hydroxylase domain proteins (PHDs) and targeted for degradation by the von Hippel–Lindau (VHL) E3 ubiquitin ligase complex (Kaelin et al., 2008). During hypoxia, hydroxylation is inhibited, resulting in the stabilization and accumulation of HIF-α subunits, which then translocate to the nucleus, dimerize with HIF-β, and bind to hypoxia response elements (HREs) in target genes, thereby enhancing their transcription (Wang et al., 1995; Majmundar et al., 2010; Zhang et al., 2014). At the whole-organism level, animals further prioritize energy expenditure by reallocating resources to sustain essential functions, such as those of the brain and heart, while reducing investment in non-essential activities like growth and reproduction. This phenomenon, observed across diverse taxa, suggests the existence of evolutionarily conserved regulatory mechanisms (Harrison et al., 2015). Despite this, the systemic factors and molecular mechanisms that orchestrate organism-wide responses to hypoxia remain poorly understood.

The zebrafish (*Danio rerio*) is a valuable vertebrate model for dissecting the genetic and endocrine mechanisms underlying somatic growth and organismal size, owing to its optical transparency, small size, rapid development, and short generation time. Previous studies in zebrafish have demonstrated that hypoxia retards developmental and growth rates by attenuating insulin-like growth factor (IGF) signaling (Kajimura et al., 2005; Kajimura and Duan, 2006; Kamei et al., 2011; Kamei, 2020). IGFs are evolutionarily ancient polypeptides structurally related to insulin. The actions of IGFs are mediated through the IGF-1 receptor (IGF-1R), a receptor tyrosine kinase. Ligand binding induces IGF-1R tyrosine phosphorylation, activating major intracellular signaling cascades such as PI3K–AKT–mTOR and RAS–MAPK–ERK pathways, which drive cell proliferation, growth, differentiation, and survival (Choi et al., 2025).

The IGF system further comprises six high-affinity IGF binding proteins (IGFBP1–6), which sequester IGFs and inhibit their interaction with IGF-1R, thereby modulating IGF signaling (Allard and Duan, 2018). Some IGFBPs also possess IGF-independent biological activities (Zhao et al., 2006; Zhong et al., 2011; Duan and Allard, 2020). The bioavailability of IGFs is tightly regulated by IGFBP proteases such as the pappalysin family members, including pregnancy-associated plasma protein-A (PAPP-A) and PAPP-A2. These enzymes cleave IGFBPs and release IGFs from the IGFBP complexes and make them available for receptor binding (Jepsen et al., 2015; Kloverpris et al., 2015). Recent findings have shown that PAPP-A-mediated IGFBP proteolysis is further modulated by stanniocalcin-1 (STC1) and stanniocalcin-2 (STC2) (Oxvig et al., 2023; Choi et al., 2025), both of which are potent PAPP-A and PAPP-A2 inhibitors (Jepsen et al., 2015; Kloverpris et al., 2015). While STC1 binds PAPP-A non-covalently, STC2 interacts covalently and irreversibly, leading to decreased IGF bioavailability and reduced IGF-1R signaling (Argente et al., 2017). Notably, human loss-of-function mutations in STC2 have been associated with increased adult height and enhanced local IGF signaling (Marouli et al., 2017; Oxvig et al., 2023).

The objective of this study is to investigate hypoxia-induced transcriptomic changes using zebrafish and to identify novel systemic factors and mechanisms regulating the trade-off between somatic growth and survival under hypoxic stress. We found that hypoxia caused major transcriptomic changes and the most prominently up-regulated pathway is hormonal activity, with *stc2a* among the top. Our genetic, physikogical, and pharmacological evidence indicates that hypoxic induction of Stc2a represents an adaptive mechanism that restricts IGF signaling, redirecting energy from somatic growth toward critical survival processes.

## Results

### Hypoxia results in major changes in hormonal signaling and metabolism

RNA-seq analysis of hypoxia-treated and control zebrafish larvae detected a total of 2411 differentially expressed genes (DEGs), including 703 up-regulated and 1708 down-regulated DEGs (Fig. 1A). The two groups are distinctly clustered in hierarchical clustering (Fig. 1B) and principal component analysis (Fig. S1A). The RNA-seq results were confirmed by qRT-PCR assays using a different set of biological samples (Fig. 1C) and similar changes were detected in 18 out of 20 genes tested (Fig. S1B).

**Figure 1.**
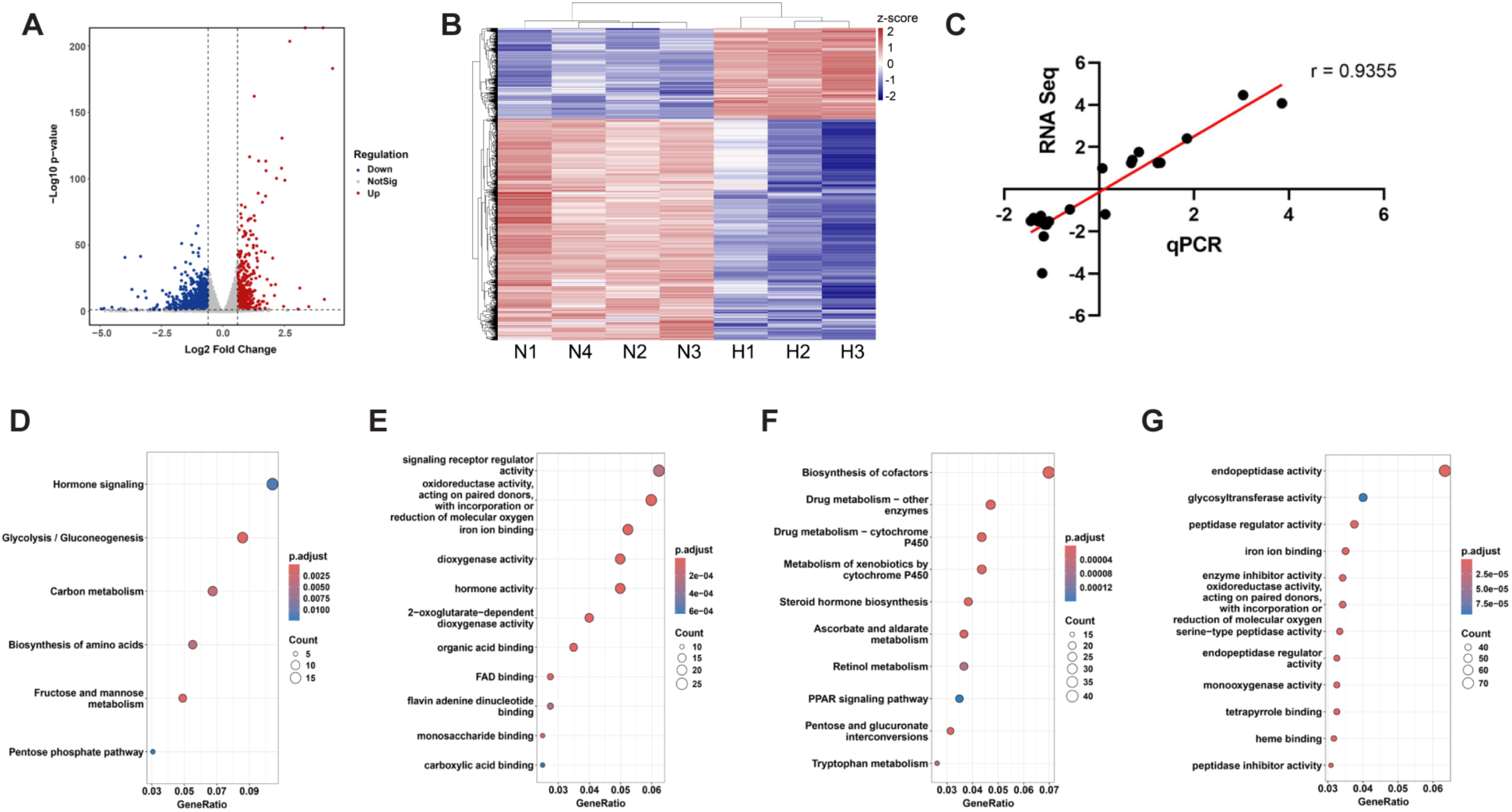
RNA-seq analysis of hypoxia-treated and normoxia control zebrafish. **(A)** Volcano plot of differentially expressed genes (DEGs) between hypoxia and normoxia groups. Significance on y-axis as -log_10_ (p-value) and effect size on x-axis as log_2_ (fold change). Dotted lines represent cutoffs of adjusted p-value < 0.05 and |log2(fold change)| > 0.6. Up-regulated genes under hypoxia are shown in red, down-regulated in blue, and non-significant genes in gray. (**B**) Hierarchical clustering of the DEGs in the normoxia (N) and hypoxia (H) groups. Rows represent individual genes, and columns represent biological replicates from hypoxia and normoxia groups. Gene expression values were normalized and scaled by row (z-scores), with red indicating higher expression and blue indicating lower expression relative to the mean. (**C**) qRT-PCR confirmation of RNA-seq data. Changes (log2) in the mRNA levels of the 20 genes measured by RNA-seq were plotted against those detected by qPCR. The line indicates the linear correlation between the results of RNA-seq and qPCR. **(D and F)** KEGG pathway enrichment for up-regulated **(D)** and down-regulated **(F)** DEGs. **(E and G)** GO molecular function enrichment analysis for up-regulated **(E)** and down-regulated **(G)** DEGs. Dot size indicates the number of genes mapped to each term, and dot color reflects the statistical significance (adjusted p-value).

KEGG analysis indicated that the up-regulated DEGs are highly enriched in the glycolysis/gluconeogenesis pathway, followed by carbon metabolism, amino acid synthesis etc. (Fig. 1D). The topmost enriched pathway, however, is hormonal signaling (Fig. 1D). Likewise, GO (Molecular Function) analysis identified "signaling receptor regulator activity" as the top enriched term in the up-regulated DEGs (Fig. 1E). "Hormone activity", largely overlapping with "signaling receptor regulator activity ", is also enriched (Fig. 1E). Other enriched GO terms are oxidoreductase activity, iron ion binding, dioxygenase activity, FAD binding etc. (Fig. 1E). The enriched down-regulated KEGG pathways include biosynthesis of co-factor, drug metabolism, cytochrome 450, steroid hormone biosynthesis etc. (Fig. 1F). The top enriched down-regulated GO terms are endopeptidase activity, glycosyltransferase activity, iron ion binding, enzyme inhibitors, endopeptidase inhibitors etc.. -1G). GO CNET plot of the up-regulated DEGs revealed the top five most significantly enriched molecular function terms are dioxygenase activity, 2-oxoglutarate-dependent dioxygenase activity, hormone activity, oxidoreductase activity acting on paired donors with incorporation or reduction of molecular oxygen, and FAD binding.

Correspondingly, the top five enriched terms among down-regulated DEGs were endopeptidase activity, peptidase regulator activity, endopeptidase regulator activity, peptidase inhibitor activity, and monooxygenase activity (Fig. S1C-D).

### Hypoxic induction of stc2a expression

We further analyzed the RNA-seq dataset by ranking the up-regulated DEGs in hormone activity according to their mRNA abundance (Fig. 2A). Among the top on the list is *stc2a*, which has been implicated in human body height regulation and IGF signaling (Argente et al., 2017; Marouli et al., 2017; Oxvig and Conover, 2023). During early development, *stc2a* mRNA levels increased gradually and reached the plateau at 4 days post fertilization (dpf) (Fig. 2B). Data mining from the Zebrahub scRNA-seq database (Lange et al., 2024) indicated that *stc2a* mRNA is detected in multiple tissues from 0 somite stages to 10 dpf, and is particularly abundant in the central nervous system, periderm, neural crest, paraxial mesoderm (Fig. 2C). In the adult stage, the highest *stc2a* mRNA levels were detected in the brain, followed by kidney, eyes, and fin. (Fig. 2D). Importantly, hypoxia increased *stc2a* mRNA levels at multiple developmental stages (Fig. 2E). The hypoxia-induced *stc2a* mRNA expression was attenuated in *hif2a*^-/-^ deficient fish (Fig. 2E), suggesting that *stc2a* expression is induced by hypoxia, likely through the action of Hif2.

**Figure 2.**
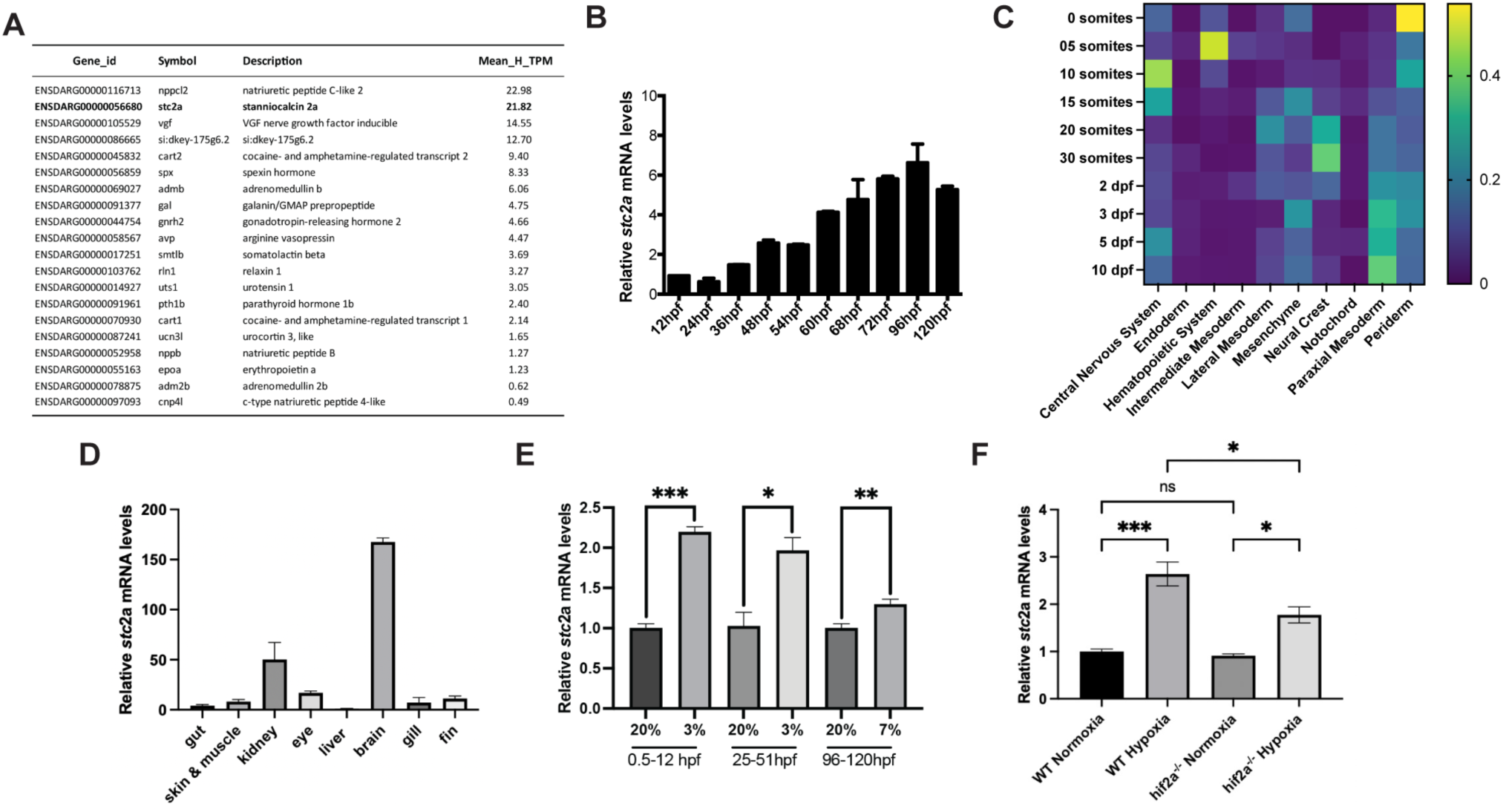
Identification of stc2a as a hypoxia-inducible gene. (**A**) List of genes enriched in the GO term hormone activity. mRNA abundance is ranked by mean transcripts per million (TPM) in the hypoxia group. (**B**) qPCR analysis results of *stc2a* mRNA expression at the indicated developmental stages. hpf, hour post fertilization. Data are shown as mean ± SEM. n = 10-15. (**C**) Relative *stc2a* mRNA expression extracted from Zebrahub scRNA-seq database. (**D**) Levels of *stc2a* mRNA in the indicated adult tissues. RNA was extracted from the indicated tissues and analyzed by RT-qPCR analysis and normalized by 18s rRNA. Data are shown as mean ± SEM. n = 3. **(E)** qPCR analysis result of *stc2a* mRNA expression. Fish were subjected to hypoxia at the indicated air O_2_ levels and periods. RNA was isolated and *stc2a* mRNA levels examined by RT-qPCR and normalized by beta-actin mRNA levels. Note: Because 3% O_2_ is lethal for advanced larvae, 7% O_2_ was used in the 96-120 hpf group. Data are shown as mean ± SEM.*p < 0.05, **p < 0.01, ***p < 0.001, t-test. n = 3-4. **(F)** Hypoxia-induced *stc2a* mRNA expression in Hif2a-deficient fish. RNA was isolated and *stc2a* mRNA levels examined by RT-qPCR, normalized by beta-actin mRNA levels.

### Loss of Stc2a increases growth of adult organ and body size

Using CRISPR-Cas9, two mutant zebrafish lines, *stc2a(Δ2+4)^-/-^*and *stc2a(Δ5)^-/-^*, were generated for functional analysis. Both are predicted to be null mutations (Fig. S2). Both mutant fish lines survived to adulthood and reproduced well under standard conditions. The gross morphology of these *stc2a^-/-^* fish was indistinguishable from their siblings (Fig. 3A). Compared with wild-type fish, however, the body length of *stc2a(Δ2+4)^-/-^*and *stc2a(Δ5)^-/-^* larvae was significantly greater (Fig. 3B). Likewise, the head-trunk angle (HTA) and somite number, two parameters of developmental speed in zebrafish (Kimmel et al., 1995), were significantly greater (Fig. 3C-D), suggesting these mutant fish develop more rapidly and grow faster. Next, *stc2a(Δ2+4)^+/-^* were intercrossed and the offsprings were grew under the same conditions for 6 months. Compared with their wild-type and heterozygous siblings, *stc2a(Δ2+4)^-/-^* fish had greater body length and body weight (Fig. 3E and 3F). Likewise, the lengths of the pelvic and dorsal spines in 1 year-old *stc2a(Δ2+4)^-/-^* fish were greater than those of the siblings (Fig. 3G and 3H). These results suggest that Stc2a negatively regulates somatic growth, adult organ and body size.

**Figure 3.**
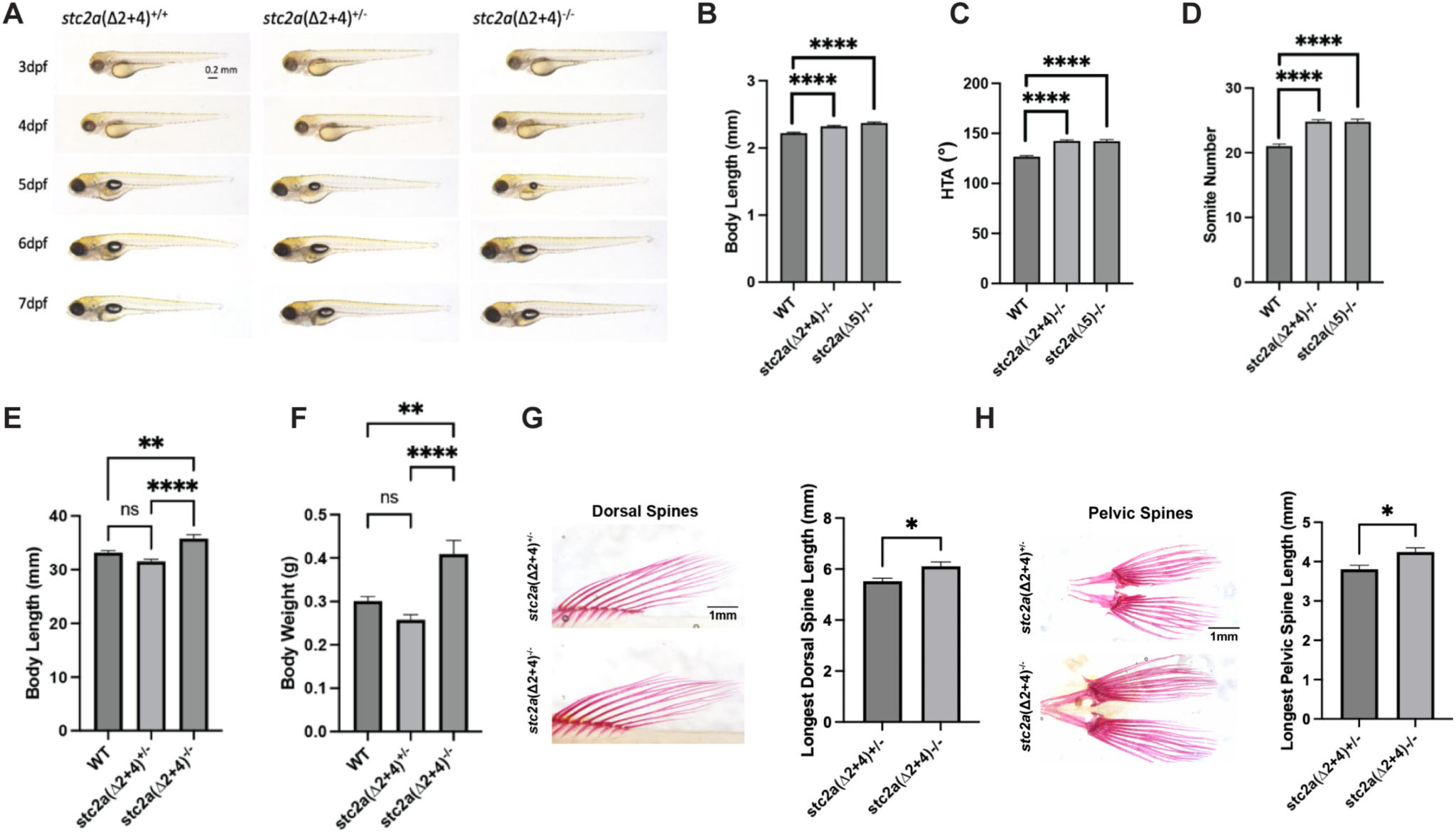
Loss of Stc2a increases developmental speed and growth rate, resulting in enlarged adult organ and body size. **(A)** Gross morphology of fish of the indicated genotypes at the indicated stages. Lateral views with anterior to the left and dorsal up. Scale bar = 0.2mm. dpf, day post fertilization. (**B-C**). Body length **(B)** and head-trunk angle (HTA) **(C)** of the indicated mutant fish and wild-type (WT) zebrafish at 51 hpf. WT, n = 76, *stc2a(*Δ*2+4)*^-/-^, n = 73, and *stc2a(*Δ*5)*^-/-^, n = 10. ****p < 0.0001 by one-way ANOVA followed by Tukey’s multiple comparisons test. **(D)** The somite number of the indicated fish line at 26 hpf. WT, n = 40, *stc2a(*Δ*2+4)*^-/-^, n = 25, and *stc2a(*Δ*5)*^-/-^, n = 14. ****p < 0.0001 by one-way ANOVA followed by Tukey’s multiple comparisons test. **(E-F)** Body length **(E)** and body weight **(F)** of the indicated genotype of 6 month-old. WT, n = 16, *stc2a(*Δ*2+4)^+/-^* n = 14, and *stc2a(*Δ*5)^-/-^* n = 15. **p < 0.01, ****p < 0.0001 by one-way ANOVA followed by Tukey’s multiple comparisons test. (**G-H**) Dorsal spine length **(G)** and pelvic spine length (**H**) of the indicated genotypes of 1 year-old. Fish were stained by alizarin red and a representative image is shown on the left and quantitative results on the right. *p < 0.05 by unpaired two-tailed t-test. *stc2a(*Δ*2+4)^+/-^* n = 6, and *stc2a(*Δ*5)^-/-^*, n = 8. In all above panels, data shown are mean ± SEM.

Since genetic deletion of Stc1a, a structurally related gene, resulted in increased ionocyte cell number, ectopic calcium deposits, kidney stone-like calcium deposits, and reduced bone mineralization (Li et al., 2021; 2023), we crossed *stc2a(*Δ*5)^-/-^* fish with *Tg* (*igfbp5a*:GFP) fish, a stable transgenic line expression EGFP in calcium transporting ionocytes or NaR cells (Liu et al., 2017; 2018), and measured the NaR cell number. There were no differences among the different genotypes (Fig. S3A-B). Alizarin red staining of the juvenile and adult *stc2a(*Δ*2+4)^-/-^*fish did not detect notable differences in calcified tissues (Fig. S3C-D), suggesting that Stc2a does not play an indispensable role in regulating ionocyte proliferation, calcium balance or bone mineralization.

### Stc2a regulates somatic growth via the Pappalysin-Igfbp-Igf signaling axis

We postulated that Stc2a may regulate somatic growth by inhibiting the pappalysin family metalloproteinase and IGF signaling activity. Commercially available antibodies against phospho-IGF1 receptor did not yield specific signal in zebrafish larvae. Western blotting of the whole body lysates did not detect major differences in phospho-Akt and phospho-Erk levels between *stc2a(*Δ*2+4)^-/-^*and wild-type fish (Fig. 4A-B). The low sensitivity of the assay and the lack of tissue resolution may obscure the results. We next tested the involvement of IGF signaling using BMS-754807, a potent IGF1R inhibitor that has been previously tested in zebrafish (Carboni et al., 2009; Kamei et al., 2011; Dai et al., 2014). If Stc2a regulates somatic growth by inhibiting IGF signaling, then inhibition of IGF signaling should reverse these phenotypes. Treatment with BMS-754807 decreased body lengths in both *stc2a(*Δ*2+4)^-/-^* fish and wild-type control fish. The magnitude of decrease, however, was significantly greater in the *stc2a(*Δ*2+4)^-/-^*fish (Fig. 4C). Likewise, ZnCl_2_, which inhibits Papp-a-mediated Igfbp degradation in human cells and in zebrafish (Tallant et al., 2006; Liu et al., 2020), caused a greater percent decrease in body length in the *stc2a(*Δ*2+4)^-/-^* larvae compared to WT siblings (Fig. 4D). To further test the role of IGF signaling, the mutant fish were treated with wortmannin, a PI3K inhibitor (Brunn et al., 1996), and U0126, a MAPK inhibitor (Favata et al., 1998). Both inhibitors decreased body length in both the wild-type and *stc2a(*Δ*2+4)^-/-^* fish. Again, the magnitude of decrease was significantly more pronounced in the *stc2a(*Δ*2+4)^-/-^* larvae (Fig. 4E-F).

**Figure 4.**
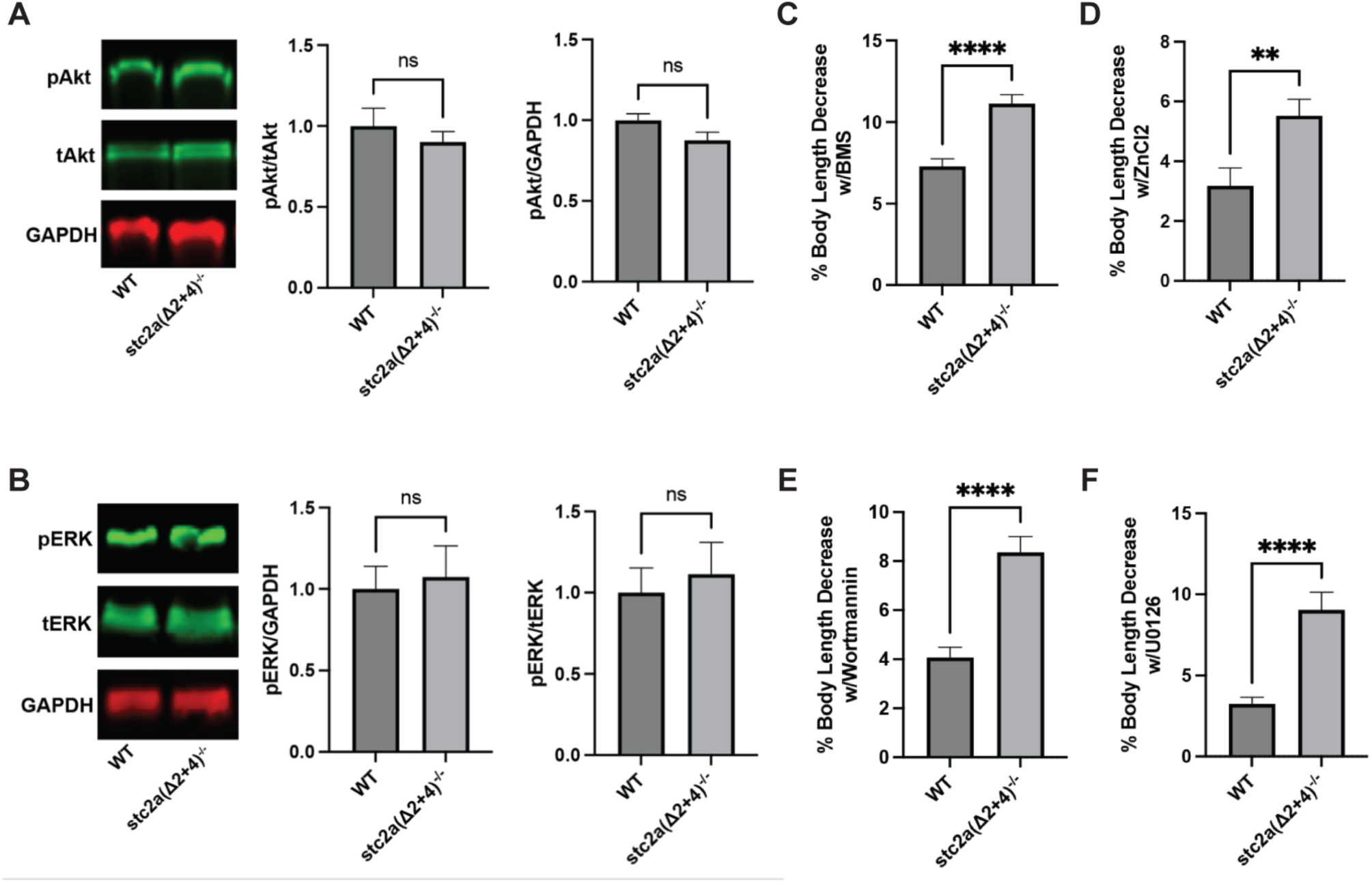
Stc2a regulates somatic growth in an IGF signaling-dependent manner. **(A)** Whole body phospho-Akt levels. Lysates of pooled wild-type (WT) and *stc2a(*Δ*2+4)*^-/-^ larvae (3 dpf) were analyzed by Western blotting using antibodies against phospho-Akt, total Akt, and GAPDH. Representative images are shown in the left panel; quantified results are shown in the middle and right panels. n = 7, ns, no significant, t -test. **(B)** The same samples described in (A) were analyzed using phospho-ERK, total ERK levels, and GAPDH. n = 7, ns, no significant, t-test. **(C)** Fish of the indicated genotypes were treated with 1.5 μM BMS-754807 or vehicle from 6-51 hpf. Body length was measured individually. Percent decrease caused by BMS treatment was calculated and shown. n = 58∼68. ***p < 0.001, t-test. (**D**) Fish of the indicated genotypes were treated with 8 μM ZnCl_2_ from 6-51 hpf. Body length was measured individually. Percent decrease caused by ZnCl_2_ was calculated and shown. n = 36∼43. *p < 0.05, t-test. (**E-F**) Fish of the indicated genotypes were treated with 0.3 μM wortmannin (E) or 8 μM U0126 or vehicle from 6-51 hpf. Body length was measured individually. Percent decrease caused by each drug was calculated and shown. n = 37-72. ****p < 0.0001, t-test. In all above panels, data shown are mean ± SEM.

### *stc2a*^-/-^ fish grew faster but had low survival rate under hypoxia

Hypoxia caused growth retardation and developmental delays in wild-type zebrafish (Fig. 6A, Kajimura et al., 2005; Kamei et al., 2011). Although hypoxia reduced the body length and HTA in *stc2a(*Δ*2+4)^-/-^* (Fig. 5A), the mutant fish still grew bigger compared to wild-type siblings under hypoxia (Fig. 5A). Likewise, while hypoxia lowered HTA values in both genotypes (Fig. 6B), the mutant fish developed faster (Fig. 5B). Treatment of mutant fish with ZnCl_2_ reduced the body length and HTA value to the levels of wild-type fish (Fig. 5C-D). We noted that mutant fish were prone to die under hypoxia and quantified their survival rate under hypoxia. They were significantly lower compared to the wild-type fish (Fig. 5E). Importantly, addition of ZnCl_2_ restored the survival rate to the levels of wild-type control groups (Fig. 5F), suggesting that pappalysin activity is required in the elevated growth and mortality in *stc2a^-/-^* mutant fish under hypoxic stress.

**Figure 5.**
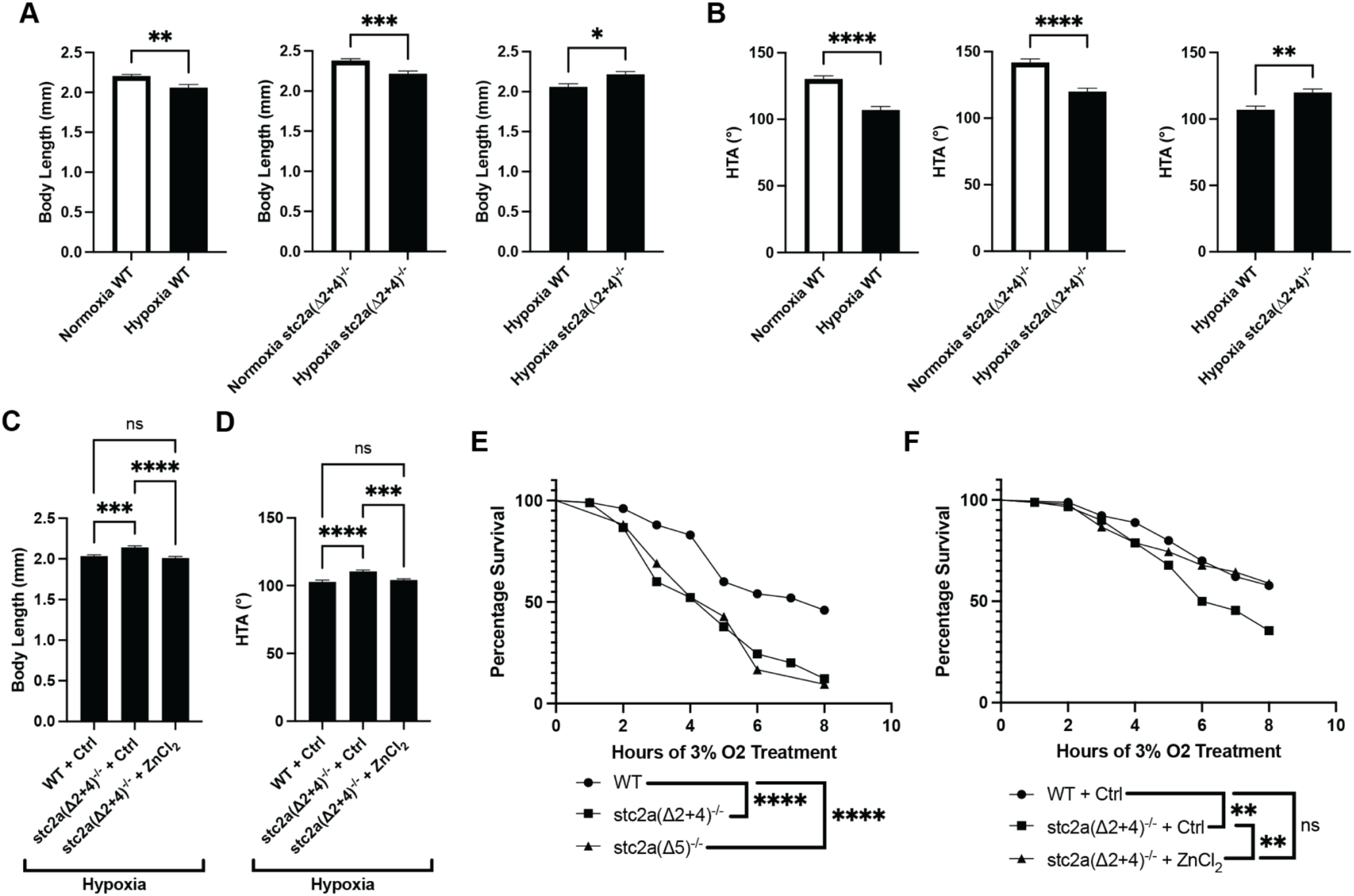
Stc2a regulates the growth-survival trade-off under hypoxia. **(A-B)** Loss of Stc2a attenuates hypoxia-induced growth retardation and developmental delay. Fish of the indicated genotypes were subjected to hypoxia (filled bar) or normoxia (open bar) from 25-51 hpf. Body length **(A)** and head-trunk angle (HTA) **(B)** were measured and shown as mean ± SEM. n = 10∼20. *p < 0.05, **p < 0.01, ***p < 0.001, ****p < 0.0001, t-test. **(C-D)** Fish of the indicated genotypes were subjected to hypoxia treatment from 25-51 hpf in the presence or absence of 8 μM ZnCl_2_. Body length (**C**) and HTA (**D**) was measured and shown as mean ± SEM. n = 58∼73. ***p < 0.001, ****p < 0.0001, One-way ANOVA with multiple comparisons. (**E**) Fish (5 dpf) of the indicated genotypes were exposed to hypoxia (3% O_2_) and survival rates were assessed and shown. ****p < 0.0001, Mantel-Cox log rank test. n = 42-100. **(F)** Fish (5 dpf) of the indicated genotypes were exposed to hypoxia (3% O_2_) in the presence or absence of 8 μM ZnCl_2_. The number of survival fish were assessed at the indicated time. ** p < 0.01, Mantel-Cox log rank test, n = 45-90.

**Figure 6.**
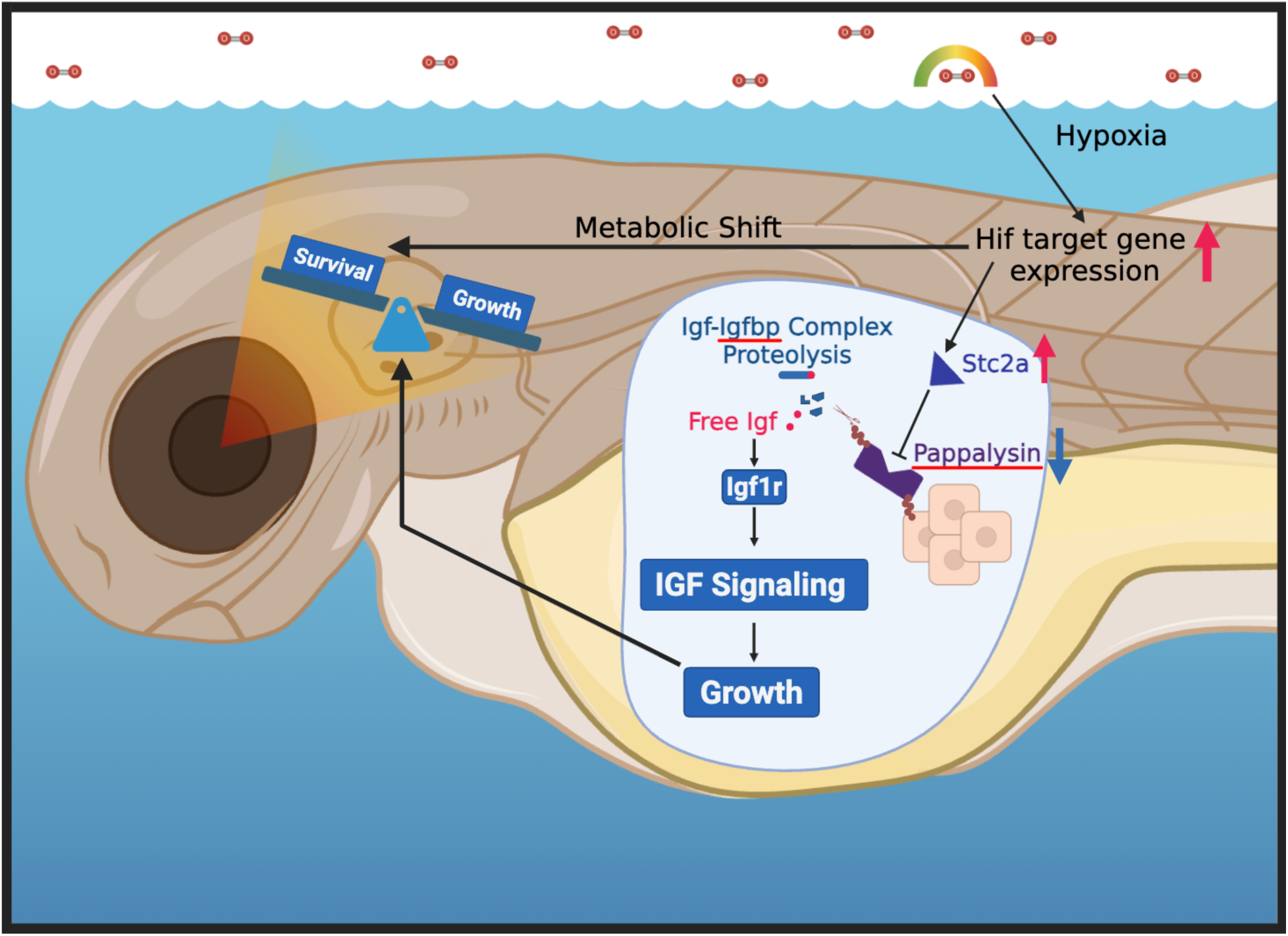
Proposed model. Hypoxia activates Hif-dependent gene expression, leading to metabolic shift towards glycolysis, protein breakdown, gluconeogenesis, and increases in hormonal signaling. An important hypoxia-induced hormonal factor is Stc2a. The increased Stc2a restricts growth and redirects toward critical survival processes by modulation of pappalysin family metalloproteinase activity and IGF signaling. Created with BioRender.com.

## Discussion

The findings made in this study suggest that Stc2a functions as an important regulator of the trade-off between somatic growth and organismal survival under hypoxic stress. We provided in vivo data showing that *stc2a* expression is highly induced by hypoxia. Loss of Stc2a increases the growth rate and developmental speed, leading to enlarged adult body size. When exposed to hypoxia, *stc2a*^-/-^ null animals exhibit accelerated growth and reduced survival. These functions are mediated by pappalysin metalloproteinase activity and IGF signaling (Fig. 6).

Stc2a is a member of the STC/Stc glycoprotein family. The first Stc protein was discovered from the Corpuscles of Stannius (CS), an endocrine organ in bony fish (Garret, 1942a; Yeung et al., 2012). Subsequent studies indicate that STC proteins are found in a wide range of species ranging from humans to eukaryotes such as Fungi, cnidarians, sponges, nematodes, and that most species have multiple STC genes (Roch and Sherwood, 2010; Schein et al., 2012). Humans, for example, have STC1 and STC2, while many teleost fish including zebrafish have 4, including *stc1a*, *stc1b*, *stc2a*, and *stc2b* (Schein et al., 2012). Previous studies have shown that loss of Stc1a, which had no effect on zebrafish somatic growth, increased the proliferation of calcium transporting ionocytes, resulting in abnormal calcium uptake, kidney stone formation, cardiac and body edema, and premature death (Li et al., 2021; 2023). The effect of Stc1a in ionocytes is mediated through its action on Papp-aa-mediated Igfbp proteolysis (Liu et al., 2020; Li et al., 2023). The functions of Stc2a and other Stc proteins, however, have not been reported. In this study, we show that genetic deletion of Stc2a increased somatic growth rate and adult body and organ size. This finding agrees with published genetic studies in mice and humans. Stc2 knockout mice exhibit increased body growth (Chang et al., 2008), while mice overexpressing human STC2 are smaller (Johnston et al., 2010). Human carriers of STC2 loss-of-functional mutation are taller than non-carriers (Marouli et al., 2017). In a genome wide association study, STC2 and its binding partners (i.e., PAPP-A and PAPP-A2) are found in loci associated with human heights (Lango Allen et al., 2010). Likewise, freshwater stickleback with different alleles with either increased or decreased Stc2a expression are associated with decreasing or increasing spine length (Roberts Kingman et al., 2021). Together, these findings suggest that STC2/Stc2a functions as a negative growth regulator in a wide range of species.

It has been suggested that STC2 inhibits somatic growth by inhibiting PAPP-A/A2- mediated IGFBP proteolysis and by reducing local IGF signaling (Oxvig and Conover, 2023). This notion is well supported by in vitro biochemical evidence. To date, there is little direct evidence showing that STC2 negatively affects body size by suppressing PAPP-a metalloproteinases and IGF signaling in vivo. Taking advantage of zebrafish larvae and the availability of *stc2a*^-/-^ fish lines, we tested the importance of Stc2a-Papp-a-IGF signaling axis on somatic growth in vivo. Our results indicate that the elevated growth observed in Stc2a deficient fish was abolished by ZnCl_2_ treatment and by pharmacological blockade of the Igf1 receptor-mediated signaling. Likewise, inhibition of PI3 kinase and MAP kinase signaling reduced the growth rate of *stc2a*^-/-^ fish to the wild-type levels. These data have provided in vivo evidence that Stc2a inhibits somatic growth by negatively inhibiting pappalysin family metalloprotease activity and IGF signaling under normoxic conditions.

A new and intriguing finding made in this study is that while loss of Stc2a increases somatic growth, it decreases organismal survival under hypoxic stress. Both actions are mediated by IGF signaling. This conclusion is supported by several lines of evidence. First, zebrafish *stc2a* is transcriptionally upregulated by hypoxia in vivo. Second, under hypoxia, while hypoxia slows down their growth rate, the mutant fish still grew faster than the wild-type fish, suggesting that stc2a deficient fish have an advantage in somatic growth under low oxygen conditions.

Meanwhile, *stc2a* mutants exhibited increased mortality under hypoxia. Both of these phenotypes were reversed by inhibiting pappalysin metalloproteinase activity. Previous reports have indicated the involvement of IGF signaling in hypoxia-induced growth retardation. Hypoxia strongly induces the expression of *igfbp1a* and *igfbp1b* mRNA in zebrafish (Kajimura et al., 2005; Kamei et al., 2008). Morpholino-based knockdown of *Igfbp1a* alleviates hypoxia-induced growth retardation by binding to IGF ligands and inhibiting their interaction with the Igf1 receptor (Kajimura et al., 2005). Likewise, the loss of *irs2b*, but not its paralog *irs2a*, blunts MAPK-activation and catch-up growth in hypoxia-treated and reoxygenated zebrafish embryos (Zasu et al., 2022). The findings made in this study reveal yet another layer of the hypoxia adaptive response, showing that a specific physiological mechanism (Stc2a-Papp-a-Igfbp) is engaged in lowering IGF signaling to prioritize energy for organismal survival over somatic growth under hypoxic stress (Fig. 6).

The notion that hypoxic induction of Stc2a inhibits IGF-stimulated somatic growth to divert resources away from growth for survival is in line with our transcriptomic analysis results. Hypoxia treatment of zebrafish caused a metabolic shift towards glycolysis and gluconeogenesis, as indicated by KEGG analysis. Among the top up-regulated KEGG pathways is glycolysis and gluconeogenesis, including *pfkpb3, hk1, pkma,* and *gpia*. Pfkfb3 is a key regulator of glycolysis and plays a crucial role for the metabolic changes seen in rapidly proliferating cancer cells, a phenomenon known as the Warburg effect (Yalcin et al., 2009; Lu et al., 2017). The hk1 gene encodes hexokinase 1, which catalyzes the first step of glycolysis. There was also a significant upregulation of *ucp3* (uncoupling protein 3), which is involved in uncoupling substrate oxidation from the ATP synthesis and reducing oxygen-dependent ATP production. (Della Guardia et al., 2024). Pkma or pyruvate kinase enzyme catalyzes the last step of glycolysis. The gpia gene encodes glucose phosphate isomerase a, which is critical in gluconeogenesis. Hypoxia treatment also resulted in a significant downregulation of *gck* (glucokinase) and *ppp1r3c2a* (Protein phosphatase 1 regulatory subunit 3C), indicating a shift away from glycogen synthesis (Taneja et al., 2024). One of the most enriched groups of down-regulated DEGs are peptidases- endopeptidases and their regulators and inhibitors. This together with the up-regulation of gluconeogenesis and amino acid synthesis is indicative of change in protein breakdown in these animals to meet the energy demand to a level that can be met by the limited oxygen supply. We speculate these changes help to divert energy from growth to maintenance and survival. Future functional studies are needed to test whether these metabolic changes are causal to the observed increase in mortality in Stc2a-deficient fish.

In conclusion, our results suggest that Stc2a limits IGF-mediated growth in favor of survival. STC2 gene has been reported to be up-regulated by hypoxia in culture human cells and mouse retina (Law et al., 2008; Law et al., 2010; Ail et al., 2022). A meta-analysis of 128 hypoxia-related human RNA-seq datasets found human STC2 upregulated in 73 datasets across varying tissue types and hypoxia severities (Bono and Hirota, 2020). Therefore, the hypoxic induction of STC2/Stc2a is likely conserved across species. Future studies will elucidate the functional importance of STC2 in the hypoxia response in mammals and humans. There may also be species differences in the HIF isoform(s) involved in the hypoxic regulation of STC2/Stc2 expression. While previous studies in human cell culture systems suggest a possible role of HIF1 in regulating STC2, we found that hypoxia caused a greater induction of *stc2a* in the Hif1a deficient fish in vivo. In comparison, the hypoxic induction of *stc2a* expression is impaired in Hif2a-deficient zebrafish, suggesting that the hypoxic induction of *stc2a* expression may be mediated by Hif2 in zebrafish. It is unclear whether the different findings are due to species difference and/or cell-type specific regulation.

### Limitations of the study

Because zebrafish larvae are tiny, several dozen were pooled for western blotting analysis. No difference was detected in phospho-Akt and phospho-Erk levels between *stc2a^-/-^*and wild-type fish, likely due to low sensitivity of Western blotting and the lack of resolution. Future studies are needed to develop more sensitive and quantitative assays to detect local IGF signaling activities. In this study, ZnCl_2_ was used as a generic pappalysin metalloproteinase inhibitor. In zebrafish, there are 3 pappalysin family members, including Papp-aa, Papp-ab, and Papp-a2 (Kjaer-Sorensen et al., 2013; 2014). The specific pappalysin metalloproteinase isoform(s) involved with the reported Stc2a action is unclear. While *papp-aa*^-/-^ mutant fish have been reported (Liu et al., 2020), these fish die prematurely. Future studies will be needed to develop conditional knockout and isoform-specific inhibitors to dissect the role(s) of Papp-aa, Papp-ab, and Papp-a2.

## Materials

Unless noted otherwise, chemical and molecular reagents were purchased from Fisher Scientific (Pittsburgh, PA, United States). Restriction enzymes were purchased from New England Biolabs (Ipswich, MA, United States) or Promega (Madison, WI, United States). TRIzol were purchased from Life Technologies (Carlsbad, CA, United States). Oligo primers were ordered from Integrated DNA Technologies (Coralville, IA,, United States). PureLink RNA Mini Kit, PureLink DNase Set, DTT, RNaseOUT Recombinant Ribonuclease Inhibitor, and M-MLV reverse transcriptase were purchased from Invitrogen (Waltham, MA, United States). Anti-GAPDH primary antibody purchased from Proteintech (Rosemont, Illinois, United States). All other primary antibodies purchased from Cell Signaling Technology (Danvers, MA, United States). Secondary antibodies purchased from LI-COR Biosciences (Lincoln, Nebraska, United States). BMS-754807 was purchased from JiHe Pharmaceutica (Beijing, China). Alizarin Red and ZnCl_2_ were purchased from Sigma (St. Louis, MO, USA). Wortmannin was purchased from Calbiochem (Gibbstown, NJ). U0126 was purchased from Selleck Chemicals (Houston, Texas).

### Experimental animals

Zebrafish were maintained following standard zebrafish husbandry guidelines (Westerfield, 2000). All experiments using zebrafish were conducted in line with guidelines approved by the Institutional Animal Care & Use Committee, University of Michigan. Embryos and larvae were raised in standard E3 medium as reported previously (Li et al., 2023). 0.003% (w/v) N-phenylthiourea (PTU) was added to the E3 medium to prevent pigmentation when required. Modified low calcium media was prepared following a previously reported protocol (Dai et al., 2014). In addition to wild-type (WT) fish, *Tg(igfbp5a:GFP)* fish (Liu et al., 2018), *hif2αb*Δ*10^-/-^* (e.g., epas1b.2, Cade et al., 2012), and *stc2a(*Δ*2+4)^-/-^* and *stc2a(*Δ*5)^-/-^*(this study) were used

### RNA sequencing and differential expression analysis

Zebrafish larvae were subjected to hypoxia (6% O₂) or normoxia (20.9% O₂, atmospheric level) from 81 to 96 hours post-fertilization (hpf). Each group comprised three to four biological replicates, with 30 larvae pooled per replicate. Total RNA was extracted using the PureLink™ RNA Mini Kit (Invitrogen) and treated with DNase using the PureLink™ DNase Set (Invitrogen). Independent RNA sample sets (n = 3∼4) were submitted for library preparation and sequencing at the Advanced Genomics Core, University of Michigan. Poly(A) RNA libraries were constructed using the NEBNext Poly(A) mRNA Magnetic Isolation Module and NEBNext UltraExpress RNA Library Prep Kit (New England Biolabs). Sequencing was performed on the Illumina NovaSeq X Plus platform, generating 89.5–104.6 million paired-end reads (151 bp) per sample. Raw sequencing reads were processed to remove low-quality bases and adapter sequences using Trimmomatic (v0.39) (Bolger et al., 2014), and read quality was evaluated before and after trimming using FastQC (v0.12.1) (Andrews, 2010). Trimmed reads were then aligned to the zebrafish reference genome (GRCz11, Ensembl release 113) using STAR (v2.7.10) (Dobin et al., 2013), with gene annotations obtained from Ensembl (Cunningham et al., 2022). DEseq2 (v1.46.0) was used to analyze the differential expression between the groups (Love et al., 2014). Genes with adjusted p-value (padj) < 0.05 and |log₂ fold change| > 0.6 were considered differentially expressed genes (DEGs). Gene Ontology (GO) and KEGG pathway enrichment analyses of DEGs were performed using the org.Dr.eg.db annotation database (v3.20.0) (Carlson) and the clusterProfiler R package (v4.14.4) (Wu et al., 2021). Enriched terms with p-value and q-value < 0.05 were considered statistically significant. Top GO terms and KEGG pathways were visualized using the dotplot function, and gene-concept networks (CNETs) were generated using the cnetplot function, both implemented in the clusterProfiler R package. All statistical analyses were conducted using R software (v4.4.1).

### Gene expression analysis by Real-time RT-qPCR and data mining

RNA was isolated from a pool of 15∼30 zebrafish larvae or from adult zebrafish tissue as reported (Schlueter et al., 2007). RNA was reverse-transcribed to cDNA using oligo-dT primers and M-MLV reverse transcriptase (Invitrogen). qPCR was performed using SYBR Green (Bio-Rad) on a StepONE PLUS real-time thermocycler (Applied Biosystems) as previously reported (Jiao et al, 2013). The expression level of a target gene transcript was normalized by 18s rRNA or β-actin mRNA levels. PCR primers were designed based on sequences as described in previous studies, the NCBI Gene database, and by NCBI Primer Blast (Ye et al., 2012; Li et al., 2021) and are listed in Table S1. The spatial expression information of stc2a mRNA was extracted from the Single-cell RNA-seq dataset from ZebraHub (https://zebrahub.org) using Scanpy (v1.11.1) (Lange et al., 2024). Expression of stc2a mRNA was extracted and averaged across tissue categories defined by annotation. Mean expression values were computed per tissue, and relative expression was calculated by normalizing to the sum of expression across all tissues. Visualization and downstream analysis were performed using Seaborn and Matplotlib.

### Generation of *stc2a^-/-^* lines by CRISPR/Cas9

Two sgRNAs targeting the *stc2a* gene were designed using CHOPCHOP (http://chopchop.cbu.uib.no/). Their sequences are: *stc2a*-gRNA1-oligo1: 5’- GCTGCTGCTCTCCGTATTGG-3’ and *stc2a*-gRNA3-oligo3: 5’-GGGTGACTCTCGTGCACATC-3’. sgRNA(30-40 ng/ul) mixed with Cas9 mRNA (200-400 ng/ul) were injected into *Tg(igfbp5a:GFP)* at the 1-cell stage. The injected F0 fish were raised to adulthood and crossed with *Tg(igfbp5a:GFP)* fish. The DNA of the F1 fish were extracted by fin clipping and analyzed by sanger sequencing. Heterozygous F1 male and female fish were crossed to generate the F2 fish.

### Genotyping

Genomic DNA, isolated from adult fin or whole larval lysate, were digested with proteinase K (60 μg/mL) in SZL buffer (50 mM KCl, 2.5 mM MgCl_2_, 10 mM Tris-HCl (pH 8.3), 0.45% NP-40, 0.45% Tween 20, 0.01% gelatine). Samples were digested at 60°C for 2 hours and at 95°C for 15 minutes. The *stc2a^-/-^* fish genotyping was performed by PCR and by direct DNA sequencing.

### Morphology and Developmental Tracking

The bright-field images of larvae zebrafish were acquired using a stereomicroscope (Leica MZ16F, Leica, Wetzlar, Germany) equipped with a QImaging QICAM camera (QImaging, Surrey, BC, Canada). Head-trunk-angle, adult zebrafish body length, body weight, and brain weight was measured following published protocols (Kamei et al., 2011). Larval body length measured from inner ear stone to tail end. Image J was used for image analysis and data quantification.

### Alizarin Red Staining

Alizarin red staining was performed as described previously (Sakata-Haga et al., 2018). The bright-field images were acquired as described above. The whole-body images were joined together with Adobe Photoshop. Dorsal and pelvic spine lengths measured by ImageJ.

### Drug Treatment

All drugs used in this study, except ZnCl_2_, were dissolved in DMSO and further diluted to desired concentrations. ZnCl_2_ was dissolved in distilled water. Zebrafish larvae were treated with drugs and vehicle as described previously (Dai et al., 2014; Liu et al., 2017). Drug solutions were changed daily.

### Hypoxia Treatment and Survival Curves

Hypoxia treatments were performed using the Invivo2 300 Hypoxia Workstation with an I-CO_2_N_2_IC advanced gas mixing system (Baker Ruskinn, Sanford, ME). Gas tanks were purchased from Cryogenic Gases (Detroit, MI). CO_2_ levels were kept constant at 1.4%. In addition to monitoring the gas values shown by this machine, an oxygen meter was utilized for a secondary reading (Sper Scientific, Scottsdale, AZ). The temperature was maintained at ∼ 28°C. Zebrafish larvae were set up in 6-well plates at a density of 15 larvae per well with 3-5 mL of E3 or drug medium. The number of dead larvae was counted hourly based on appearance and motor response to stimulus.

### Western Blotting

Zebrafish larvae (30-40 per group) were homogenized in RIPA buffer (50 mM Tris-HCl, 150 mM NaCl, 1 mM EGTA, 0.1% Triton X-100, 0.5% sodium deoxycholate, 0.1% SDS, pH 7.4) containing a cocktail of protease inhibitors and phosphatase inhibitors on ice for 30 seconds. Zebrafish homogenates were centrifuged at 13,000 rpm, 4°C, for 20 minutes. The protein concentration of the supernatant was measured by Bradford assay (Bio-rad) and normalized. 2X urea loading buffer (150 mM Tris pH 6.8, 6 M Urea, 6% SDS, 40% glycerol, 100 mM DTT, 0.1% Bromophenol blue). Samples were heated at 95°C for 5 minutes and subjected to 12% SDS-PAGE gels, and transferred to a nitrocellulose membrane for western blot analysis. After incubation with primary and secondary antibodies, membranes were scanned using the Odyssey CLx imaging system (LI-COR). The ratios of phosphorylated to total protein were calculated after the images had been analyzed. The primary antibodies were mouse anti-GAPDH (1:4000, Proteintech), rabbit anti-Akt (1:1000, Cell Signaling Technology), rabbit anti-P-Akt (S473) (D9E) XP® (1:2000, Cell Signaling Technology), rabbit anti-p44/42 MAPK (1:2000, Cell Signaling Technology), rabbit anti-Phospho-p44/42 MAPK (D13.14.4E) XP® (1:2000, CellSignaling Technology). The secondary antibodies, goat anti-mouse IRDye 680LT and goat anti-rabbit IRDye 800CW were purchased from LI-COR Biosciences and used at 1:10,000 dilution.

### Statistical Analysis

Statistical analysis was performed using GraphPad Prism 9 software (GraphPad Software, Inc., San Diego, CA). Values are shown as means ± SEM. Statistical significance between experimental groups was performed using unpaired two-tailed t-test, one-way ANOVA followed by Tukey’s multiple comparison test, or two-way ANOVA with multiple comparisons. Differences between the experimental groups in survival curves were analyzed using the Mantel-Cox log-rank test. Statistical significances were accepted at *p < 0.05, **p<0.01, ***p < 0.001, ****p < 0.0001.

---

**Figure S1.**
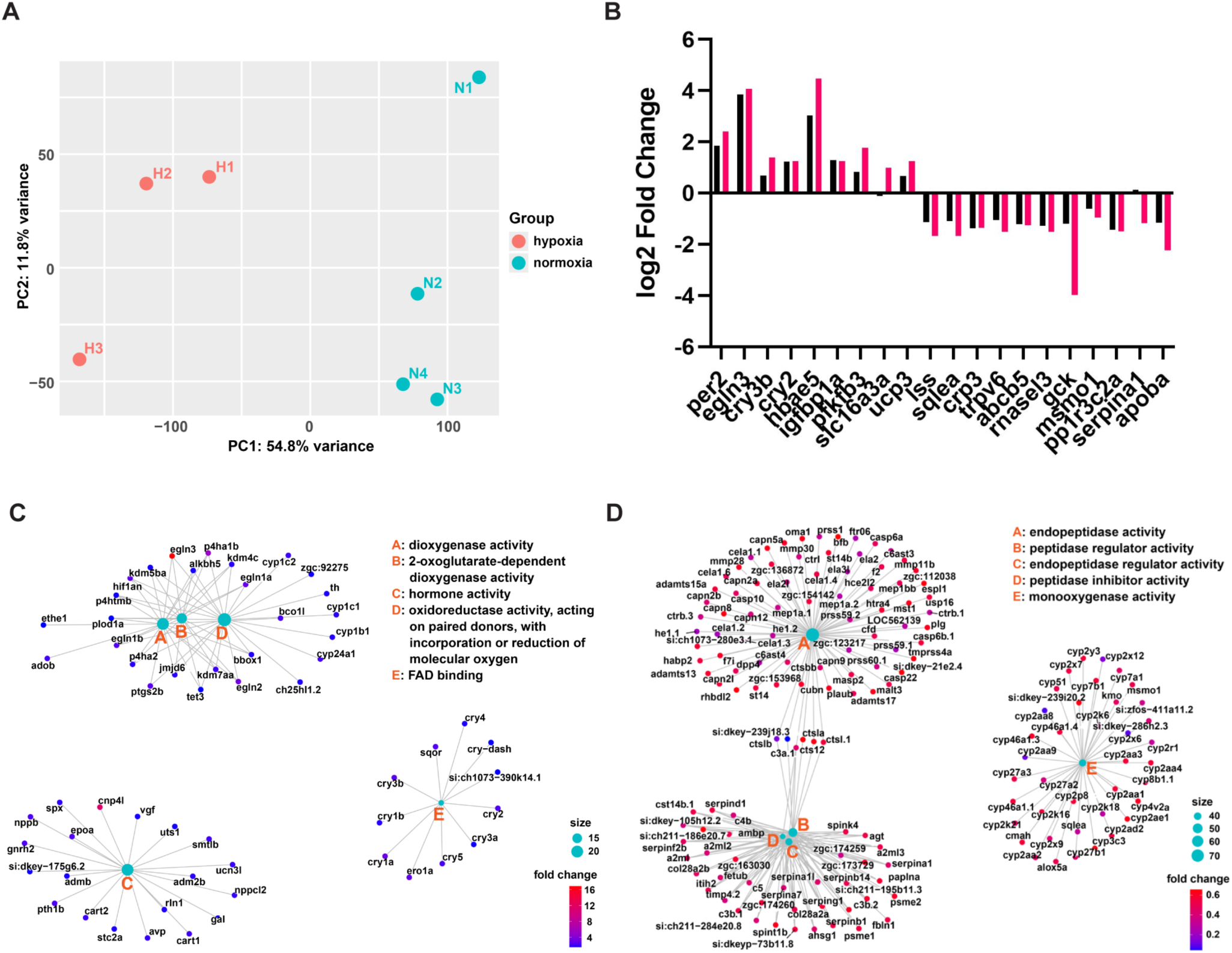
(A) Principal component analysis (PCA) plot of normoxia (N) and hypoxia (H) groups. The plot shows the first two principal components (PC1 and PC2) of normalized gene expression data, illustrating the overall variance and separation between normoxia and hypoxia groups**. (B)** The mRNA levels of the indicated genes were measured by qPCR (blue) or RNA-seq (red) and shown as the ratio between the hypoxia groups and the normoxia groups. n = 2-4. (**C and D**) Gene-concept networks (CNETs) of differentially expressed genes. CNETs show the top five significantly enriched GO molecular function terms for (**C**) up-regulated and (**D**) down-regulated DEGs. Small nodes represent genes associated with each GO term, with node color indicating fold change, whereas large cyan nodes represent the GO terms.

**Figure S2.**
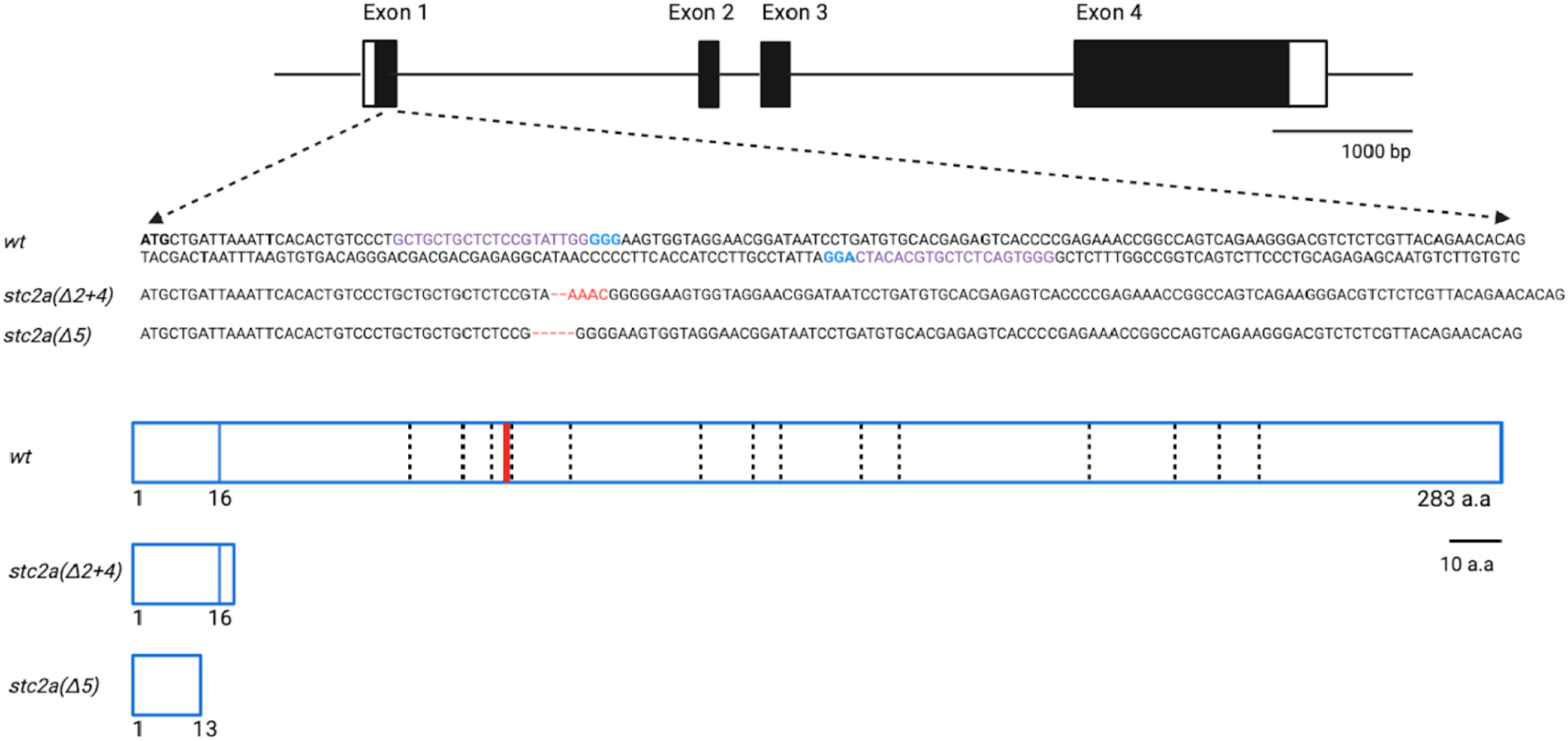
Schematic diagram of zebrafish *stc2a* gene and the engineered mutants. Boxes represent exons and lines represent introns. Filled boxes represent protein coding regions and open boxes represent untranslated regions. The PAM motif is represented in blue color. Dashed lines indicate conserved cysteine residues. Blue boxes indicate N-glycosylation sites.

**Figure S3.**
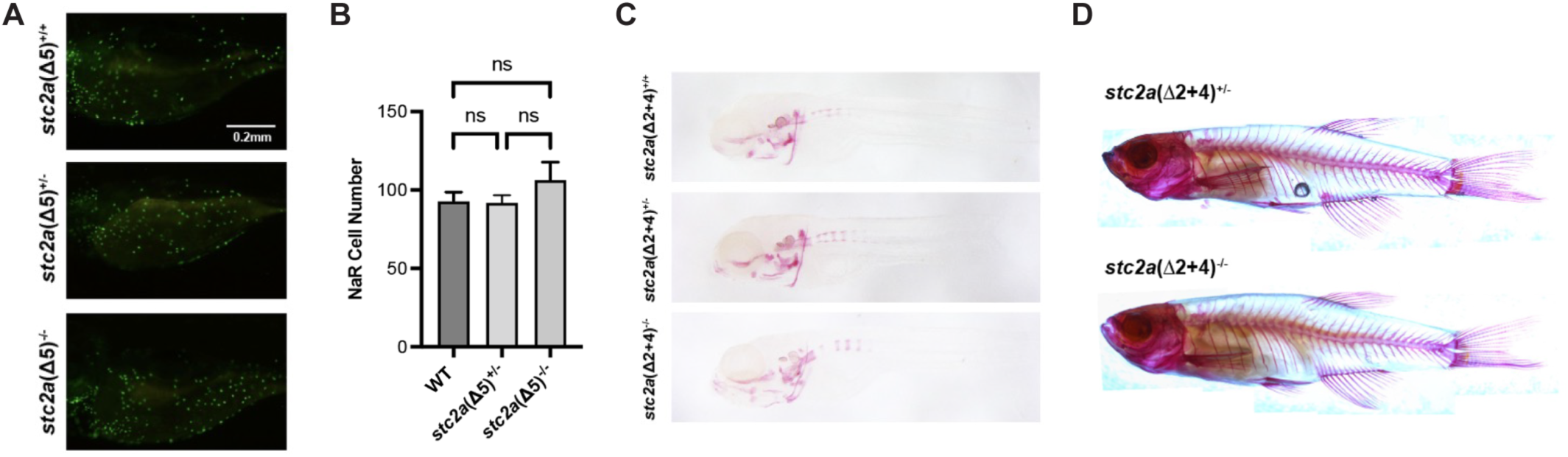
Loss of Stc2a does not change ionocyte cell proliferation or bone mineralization. **(A-B)** NaR cell number of the indicated genotype fish in the Tg(igfbp5a:GFP) background were measured at 5 dpf. Representative images of GFP-expressing NaR cells are shown in **(A)** and quantified results in **(B).** Data are shown as mean ± SEM. n = 5-18, ns, not statistically significant, one-way ANOVA followed by Tukey’s multiple comparisons test. (**C**) Representative images of 7 dpf fish of the indicated genotypes stained by Alizarin red. (**D**) Representative images of 1 year-old adult fish stained by Alizarin red.

